# Selectivity filter instability dominates the low intrinsic activity of the TWIK-1 K2P K^+^ Channel

**DOI:** 10.1101/735704

**Authors:** Ehsan Nematian-Ardestani, M. Firdaus Abd-Wahab, Franck C. Chatelain, Han Sun, Marcus Schewe, Thomas Baukrowitz, Stephen J Tucker

**Author notes:** To whom correspondence should be addressed: Stephen J. Tucker, Clarendon Laboratory, Department of Physics, University of Oxford, OX1 3PU, United Kingdom.

## Abstract

Two-pore domain (K2P) K^+^ channels have many important physiological functions. However, the functional properties of the TWIK-1 (K2P1.1/*KCNK1*) K2P channel remain poorly characterized because heterologous expression of this ion channel yields only very low levels of functional activity. Several underlying reasons have been proposed, including TWIK-1 retention in intracellular organelles, inhibition by post-translational sumoylation, a hydrophobic barrier within the pore, and a low open probability of the selectivity filter (SF) gate. By evaluating these various potential mechanisms, we found that the latter dominates the low intrinsic functional activity of TWIK-1. Investigating the underlying mechanism, we observed that the low activity of the SF gate appears to arise from the inefficiency of K^+^ in stabilizing an active (*i.e.* conductive) SF conformation. In contrast, other permeant ion species, such as Rb^+^, NH_4_^+^, and Cs^+^, strongly promoted a pH-dependent activated conformation. Furthermore, many K2P channels are activated by membrane depolarization via a SF-mediated gating mechanism, but we found here that only very strong, non-physiological depolarization produces voltage-dependent activation of heterologously expressed TWIK-1. Remarkably, we also observed that TWIK-1 Rb^+^ currents are potently inhibited by intracellular K^+^ (IC_50_ = 2.8 mM). We conclude that TWIK-1 displays unique SF gating properties among the family of K2P channels. In particular, the apparent instability of the conductive conformation of the TWIK-1 SF in the presence of K^+^ appears to dominate the low levels of intrinsic functional activity observed when the channel is expressed at the cell surface.

---

K2P K^+^ channels represent a structurally unique family of channels involved in diverse physiological functions such as cell-volume regulation, apoptosis, vasodilation, central chemosensitivity, neuronal excitability and the perception of pain. They also act as a major target for volatile anesthetics and represent attractive therapeutic targets for the treatment of a wide variety of cardiovascular and neurological disorders (1-3).

The first mammalian member of this family to be identified was the TWIK-1 (K2P1/*KCNK1*) channel (4). This channel has subsequently been implicated in a number of important physiological roles in the heart and the brain (5-7). However, understanding the biophysical, pharmacological and functional properties of TWIK-1 has proven difficult because of the extremely low levels of functional activity observed when the wild-type (WT) channel is recombinantly expressed in heterologous systems such as *Xenopus* oocytes or cultured mammalian cells (e.g. HEK293) that are typically used for electrophysiological recordings of ion channels.

The precise reasons underlying these low levels of functional activity remain unclear. Originally, it was suggested that post-translational modification (sumoylation) of a lysine residue (Lys^274^) within the proximal C-terminus was responsible for its apparent lack of functional activity (8). The structure of TWIK-1 revealed Lys^274^ to be located within an amphipathic interfacial ‘C-helix’ structure kinked parallel to the membrane close to the intracellular entrance to the pore (9). However, the mechanisms underlying the increased activity seen with mutations at Lys^274^ within the C-helix remain controversial (10).

A subsequent study of wild-type TWIK-1 channels revealed they are actively and constitutively internalized from the cell surface, and that mutation of an endocytic motif within the intracellular C-terminal domain can reduce this internalization to increase channel levels within the plasma membrane (11). However, even when engineered to reside on the cell surface in greater number, these ‘WT’ TWIK-1 channels still exhibit very low levels of functional activity compared to many other members of the K2P channel family.

Exploiting this approach to stabilize TWIK-1 within the plasma membrane, it was also found that larger TWIK-1 currents can be produced by mutation of hydrophobic residues that form a ‘hydrophobic cuff’ within the inner pore of the channel (e.g. Leu^146^). This cuff creates a hydrophobic barrier that restricts hydration of the inner cavity below the selectivity filter (12-15). However, the precise role of this barrier also remains unclear because in addition to its effect on pore hydration, mutation of residues that form this barrier may also indirectly affect other aspects of K2P channel gating (16). Overall, detailed functional characterization of non-mutant TWIK-1 channels therefore remains challenging.

Another unusual structural feature of the TWIK-1 channel is its highly asymmetric and non-canonical selectivity filter (SF). By nature of their dimeric architecture, all K2P channels exhibit a two-fold symmetry and have a SF formed from non-identical sequences, but for TWIK-1 there are also major variations in the canonical ‘TXGY/FG’ sequence with ‘TTGYG’ in the first P-loop and ‘TIGLG’ in the second. Furthermore, the channel has been shown to exhibit a dynamic ion selectivity with reduced K^+^>Na^+^ selectivity that can be modulated by mutations either within or close to the first P-loop (15,17,18). Interestingly, this dynamic ion selectivity is thought to be relevant for the regulation of cardiac excitability under conditions of hypokalemia (5,7).

In most other K2P channels, the SF also represents the main site for channel gating and the origin of their voltage sensitivity (19-22). In a previous study, we demonstrated this voltage-dependent gating originates from the movement of ions into the high electric field of the SF to generate a one-way “check valve” within the filter where the outward movement of K^+^ induces filter opening, whereas inward movement promotes channel closure. The gating process was also highly dependent upon the nature of the permeant ion (23). However, at that time, the relevance of this filter gating mechanism to TWIK-1 was not examined because of its apparent linear current-voltage behavior within the physiological voltage range. Nevertheless, it had previously been shown that macroscopic TWIK-1 currents could be recorded when Rb^+^ was used as the permeant ion (12,17), thus suggesting the presence of some type of filter-gating mechanism similar to that seen in other K2P channels.

In this study, we therefore sought to understand the extent to which these different possible mechanisms contribute to the low levels of functional activity associated with TWIK-1. We found that TWIK-1 does indeed possess a filter-gate that displays markedly different characteristics to most other K2P channels and appears the principal reason why TWIK-1 produces very low levels of channel activity when heterologously expressed.

## Results

### Role of the C-Helix

To begin to evaluate the effect of these different mechanisms, we first examined the role of the C-helix. A lysine residue within the C-helix (Lys^274^) has been proposed to be a site for post-translation sumoylation that silences TWIK-1 activity (8). However, the C-helix also contains a number of other charged residues. We therefore conducted a scanning mutagenesis of the C-helix and compared the levels of activity produced when these mutant channels were expressed in *Xenopus* oocytes.

Constitutive endocytosis has previously been reported to reduce the levels of TWIK-1 within the plasma membrane (11,15). Therefore, to permit the measurement of larger currents, all mutations were created in a background construct where the di-isoleucine retention motif was mutated (I293A/I294A). This site is located within the cytoplasmic domain and consistent with many previous studies of TWIK-1, this trafficking mutant (hereafter referred to as TWIK-1*) is used throughout this study to measure channel activity.

As previously reported (12), we found that mutation of K274 in TWIK-1* caused an increase in whole-cell currents. However, although we found that most other mutations in the C-helix had little effect, mutation of two other charged residues (Arg^277^ and Lys^278^) also increased the level of whole-cell current activity (**Fig. 1A and Supplemental Fig. S1**). Interestingly, when mapped on to the structure of the C-helix, these three charged residues were observed to face towards the entrance of the inner cavity. By contrast, mutation of adjacent charged residues (e.g. Glu^272^ and Lys^275^) that face away from the pore, had no such effect (**Fig. 1A and B**).

**Figure 1.**
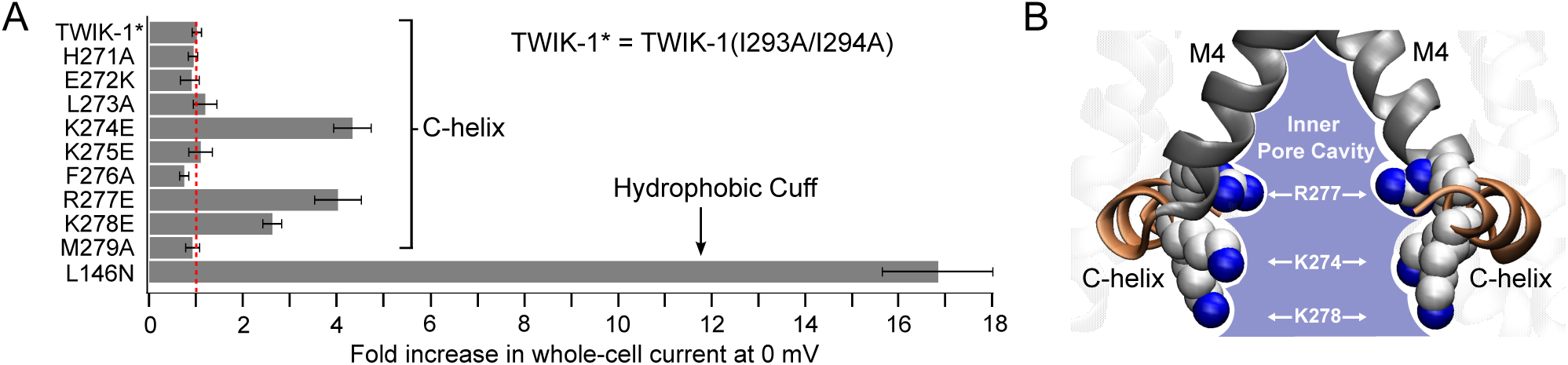
Role of charged residues within the C-helix. **A)** Fold-increase in activity of TWIK-1 mutations in the C-helix in comparison to a mutation within the hydrophobic cuff (L146N). Maximum whole cell currents were recorded at 0 mV from channels expressed in *Xenopus* oocytes. All mutations were introduced into the TWIK-1* construct that exhibits greater stability within the plasma membrane. The dotted red line indicates the level of current recorded for WT TWIK-1* (0.8 ± 0.1 µA at 0 mV, n≥18; see also supplemental Fig. S1). **B)** The three activatory mutations within the C-helix are positively charged and point in towards the inner mouth of the pore. Mutation of other charged residues that point away do not increase currents.

The precise mechanisms responsible for the increased activity of these pore-facing mutations remains unclear and likely to involve multiple factors. However, the fact these three pore-facing mutants were all able to increase macroscopic current levels to the same levels as mutation of Lys^274^ strongly suggests that the role of Lys^274^ in ‘silencing’ this channel may not be as unique as previously thought.

Importantly, and consistent with previous reports, the increase in currents seen with these C-helix mutants is also relatively small in comparison to the much bigger effect of mutations within the hydrophobic cuff of the inner pore, e.g. the L146N mutation in M2 (**Fig. 1A**). Furthermore, we have also shown that both WT TWIK-1* and L146N mutant channels produce even larger macroscopic currents when Rb^+^ is used as the permeant ion (12). This indicates that the low levels of TWIK-1* activity typically observed when the channel is expressed can only partly be accounted for by the C-helix, and the intrinsic biophysical properties of the SF are likely to play a more dominant role in this process.

### Functional properties of TWIK-1 currents using Rb^+^ as the permeant ion

Our previous studies identified an ion flux-gating mechanism within the SF of many different subfamilies of K2P channels (23), but its contribution to TWIK-1 gating remains unclear. We therefore next examined how the SF gate contributes to the unusual functional properties of TWIK-1.

When K^+^ is used as the permeant ion with either WT TWIK-1 or TWIK-1*, relatively little current can normally be recorded above background levels within a physiological voltage range (i.e. -100 to +100 mV) (4,23). However, **Fig. 2A** shows that in excised patches, strong depolarization in excess of +100 mV begins to elicit measurable K^+^ currents compared to uninjected controls. These currents display similar time- and voltage-dependent activation properties observed in other K2P channels such as TREK-1 and TRAAK. However, unlike these other K2P channels, the threshold for this activation is shifted to >100 mV highlighting the unusual flux-gating properties of the SF gate in TWIK-1*.

**Figure 2.**
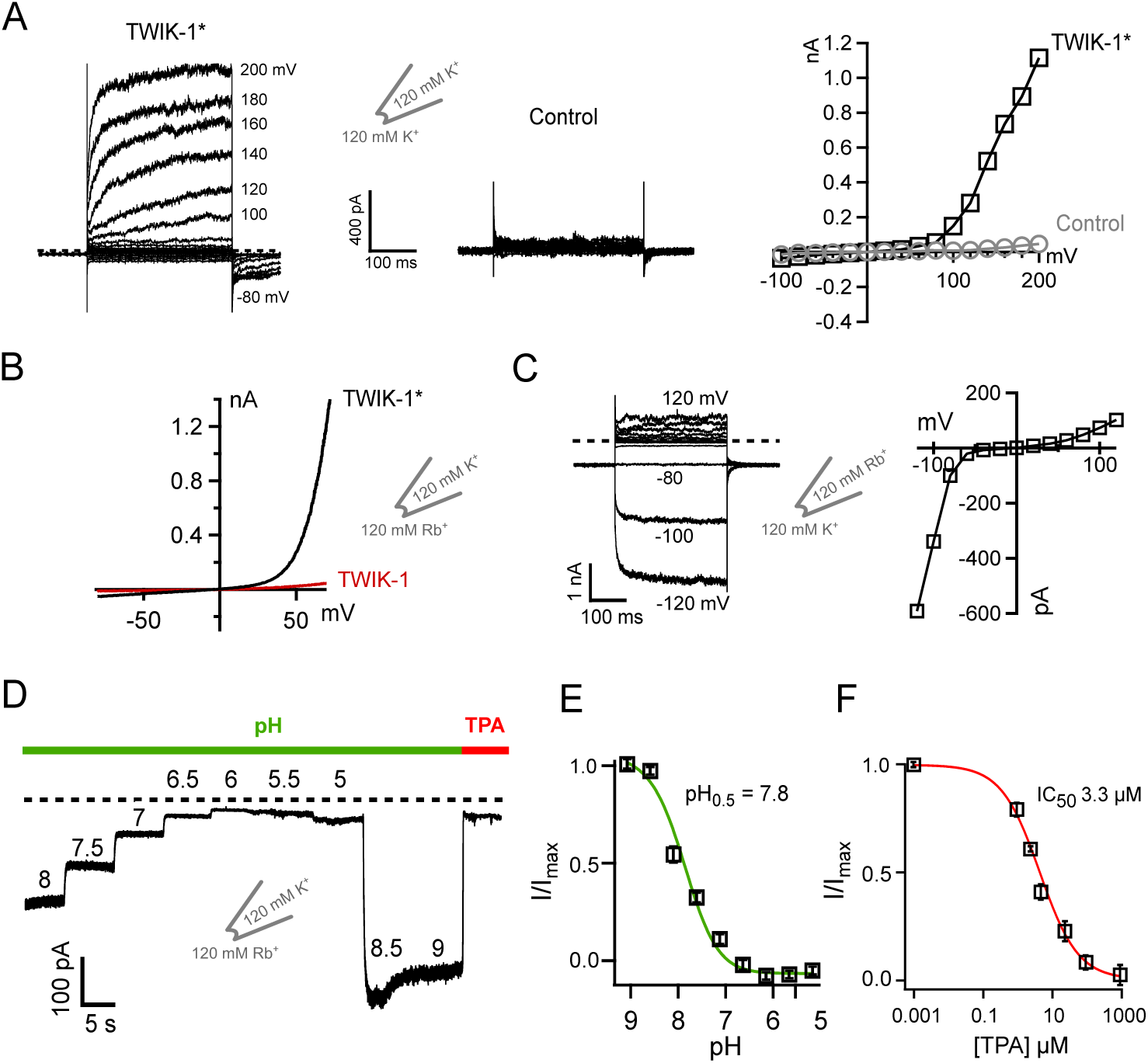
Functional characterization of TWIK-1* channels in excised patches. **A)** Example of K^+^ currents recorded from TWIK-1* in giant excised patches in response to voltage steps from -100 mV to + 200 mV. Note the supra-physiological voltages (>100mV) required to observe voltage-dependent activation. At +140 mV TWIK-1* currents averaged 0.49 ± 0.1 nA, in comparison to 0.01 ± 0.005 nA for uninjected controls, *n*≥6. **B**) Comparison of the currents recorded in giant excised patches from oocytes expressing wild-type TWIK-1, or the TWIK-1* trafficking mutant, I293A/I294A. Currents were recorded in response to a voltage ramp from -70 mV to + 70 mV. Patches have 120 mM Rb^+^ on the inside with 120 mM K^+^ in the pipette solution. **C)** Left: activation of TWIK-1* currents when K^+^ is replaced with Rb^+^ on the extracellular side. Currents recorded in excised patches from oocytes in response to a series of voltage steps from +120 to -120 mV with 120 mM Rb^+^ in the pipette and 120 mM intracellular K^+^. Right: a current voltage plot from this example recording. **D)** Intracellular pH sensitivity of TWIK-1* Rb^+^ currents. **E)** Normalized TWIK-1* Rb^+^ currents plotted against intracellular pH, and **F)** plotted against TPA concentration.

When Rb^+^ is used as the permeant ion in TREK and TRAAK channels, it activates the SF ion-flux gating mechanism within the SF by stabilization of its active configuration (23,24). Consistent with this, we also observe that Rb^+^ is capable of producing large currents in inside-out macropatches excised from *Xenopus* oocytes expressing WT TWIK-1*. These currents were also activated at less depolarized potentials than are required to generate K^+^ currents (**Fig. 2B**). This suggests that K^+^ is much less efficient than Rb^+^ in stabilizing the SF in an actively conductive configuration. **Figure 2B** also highlights the relative importance of the endocytotic retention motif because when the retention motif is present (i.e. WT TWIK-1) then the currents are markedly smaller, even with Rb^+^ as the permeant ion. The ability to record robust Rb^+^ currents from WT TWIK-1* therefore provides an opportunity for a more detailed biophysical investigation of the intrinsic gating properties of this unusual K2P channel.

In the excised macropatch recordings described above, Rb^+^ is applied intracellularly and its activatory effect is consistent with the intracellular effects of Rb^+^ seen for many other K2P channels. However, consistent with previous reports (17), we also observed that this activatory effect occurs extracellularly, i.e. when Rb^+^ was included in the pipette solution (**Fig. 2C**). This is in contrast to many other K2P channels (e.g. TREK-1) where Rb^+^ activation only works from the intracellular side (23). This provides further evidence that the TWIK-1 SF gating mechanism is different to most other members of the K2P channel family.

TREK/TRAAK channel activity can also be regulated by intracellular H^+^ and so we examined whether intracellular pH also regulates TWIK-1* activity. **Figs. 2D and E** show that the robust Rb^+^ currents recorded from TWIK-1* in excised patches are also sensitive to inhibition by intracellular H^+^ (pH_0.5_ ∼ 7.8). Thus, to optimize currents all experiments were subsequently performed using an intracellular pH of 8.0. Furthermore, to distinguish TWIK-1* currents from leak, we quantified the effects of intracellular tetrapentylammonium (TPA) as a channel blocker (**Fig. 2F**). We found that intracellular TPA blocked TWIK-1* Rb^+^ currents with an even higher affinity (IC_50_ = 3.3 ± 0.1 µM) than typically observed for most other K2P channels (21,25). These results therefore established an experimental framework for the measurement of robust TWIK-1 channel activity.

### Stabilization of the SF gate by different permeant ions

Several other monovalent cations have also been shown to activate the ion-flux gating mechanism in other K2P channels, with Rb^+^ and NH_4_^+^ typically being the most potent, and Na^+^ the least (23). **Figs. 3A-C and supplemental Fig S2** show a similar pattern of activation for WT TWIK-1*. But, in marked contrast to other K2Ps where Cs^+^ acts as a permeant blocker, Cs^+^ appears to activate the SF gate in TWIK-1* more effectively than K^+^. Furthermore, despite the levels of activation induced by intracellular NH_4_^+^ and Rb^+^, no tail currents could be observed upon repolarization (**Fig. 3A & D**). This is in marked contrast to many other K2P channels where, under similar conditions, large tail currents can be observed upon repolarization (23).

**Figure 3.**
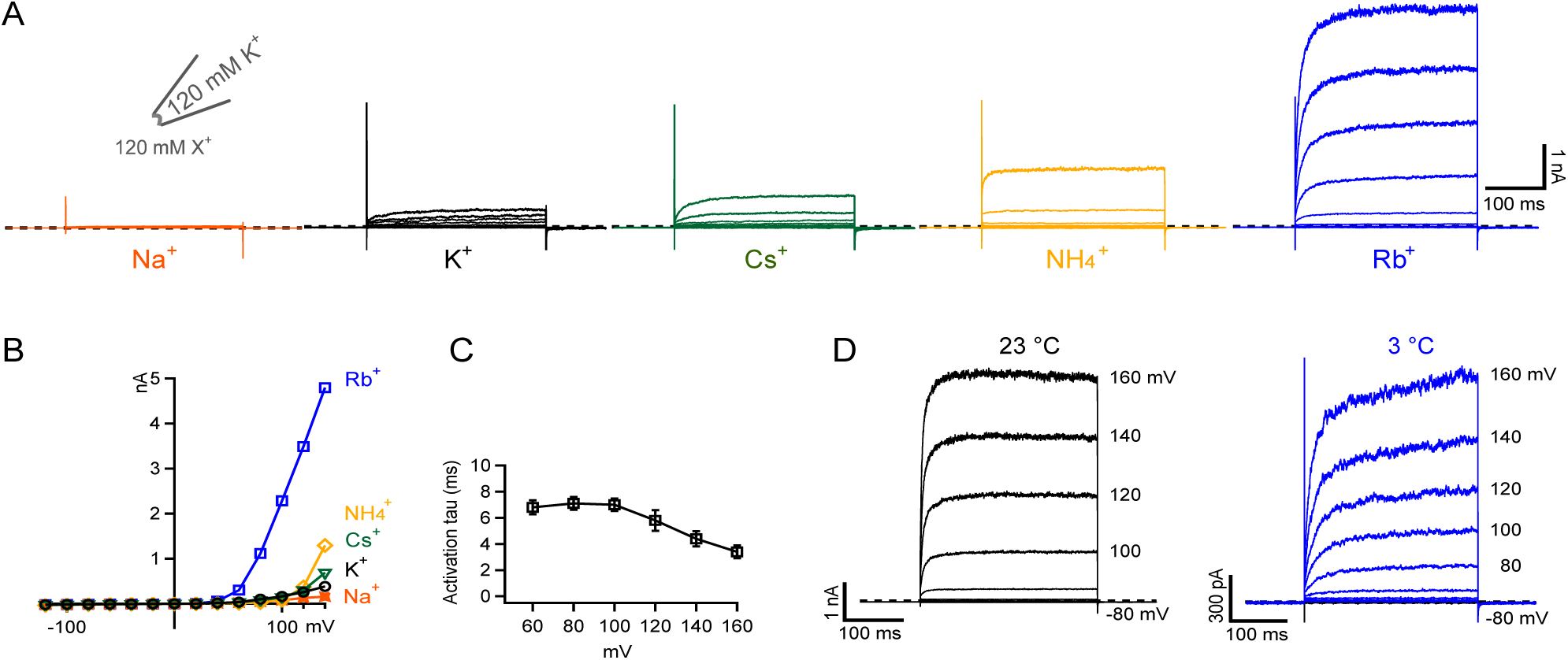
Role of different permeant ions. **A)** Examples of currents recorded in a single giant excised inside-out patch from an oocyte expressing TWIK-1* in response to a series of voltage steps from -120 to +140 mV. 120 mM of each of the indicated ions was exchanged on the intracellular side. Traces are ordered based on the potency of activation by each ion. **B)** Current voltage plots for each of these different ions. **C)** The rate of activation by Rb^+^ appears to be independent of the applied voltage. **D)** Comparison of TWIK-1* Rb^+^ activation at room temperature and at 3°C. Despite the obvious decease in the rate of activation, still no tail currents are observed at 3°C (see also supplemental Fig. S2B).

The activation of TWIK-1* by extracellular Rb^+^ (**Fig. 2C**) rules out the possibility that the channel cannot conduct extracellular ions into the cell, but to examine the apparent absence of tail currents further, we repeated these measurements at 3 °C to slow down channel gating kinetics. As a control for this effect, we used a mutant version of TREK-2 (26) where the normally large tail currents have become so fast as to appear absent at 23°, but reducing the temperature slows the kinetics of activation and inactivation in this version of TREK-2 (**supplemental Fig S2**). However, for TWIK-1*, although the rate of channel activation also decreased at 3 °C, the inactivation kinetics of TWIK-1 were still unable to be resolved (**Fig. 3D** and **supplemental Fig S2)**.

### The SF is highly unstable in the presence of K^+^

Based upon these observations, we hypothesized that when the SF is occupied by K^+^ it may be extremely unstable resulting in almost instantaneous inactivation when Rb^+^ is replaced by K^+^. To investigate this further, Rb^+^ was used as the permeant ion and increasing concentrations of K^+^ were added. Intriguingly, we found that very low concentrations of K^+^ strongly inhibited these Rb^+^ currents (**Fig. 4A**). A full dose-response curve for this effect revealed an IC_50_ for K^+^ of 2.8 mM (**Fig. 4A**). In marked contrast, the TREK-1 K2P channel, showed comparatively weaker K^+^-dependent inhibition of Rb^+^ currents (IC_50_ of ∼ 68 mM). To test whether this observed inhibitory effect of K^+^ on WT* TWIK-1 currents is specific, we compared the effect of several other monovalent cations. We found that K^+^ exhibited the strongest inhibitory effect and Cs^+^ the weakest (**Fig. 4B**). This also appears to be in agreement with the relative activatory effect of these different ions shown in **Fig. 3A**.

**Figure 4.**
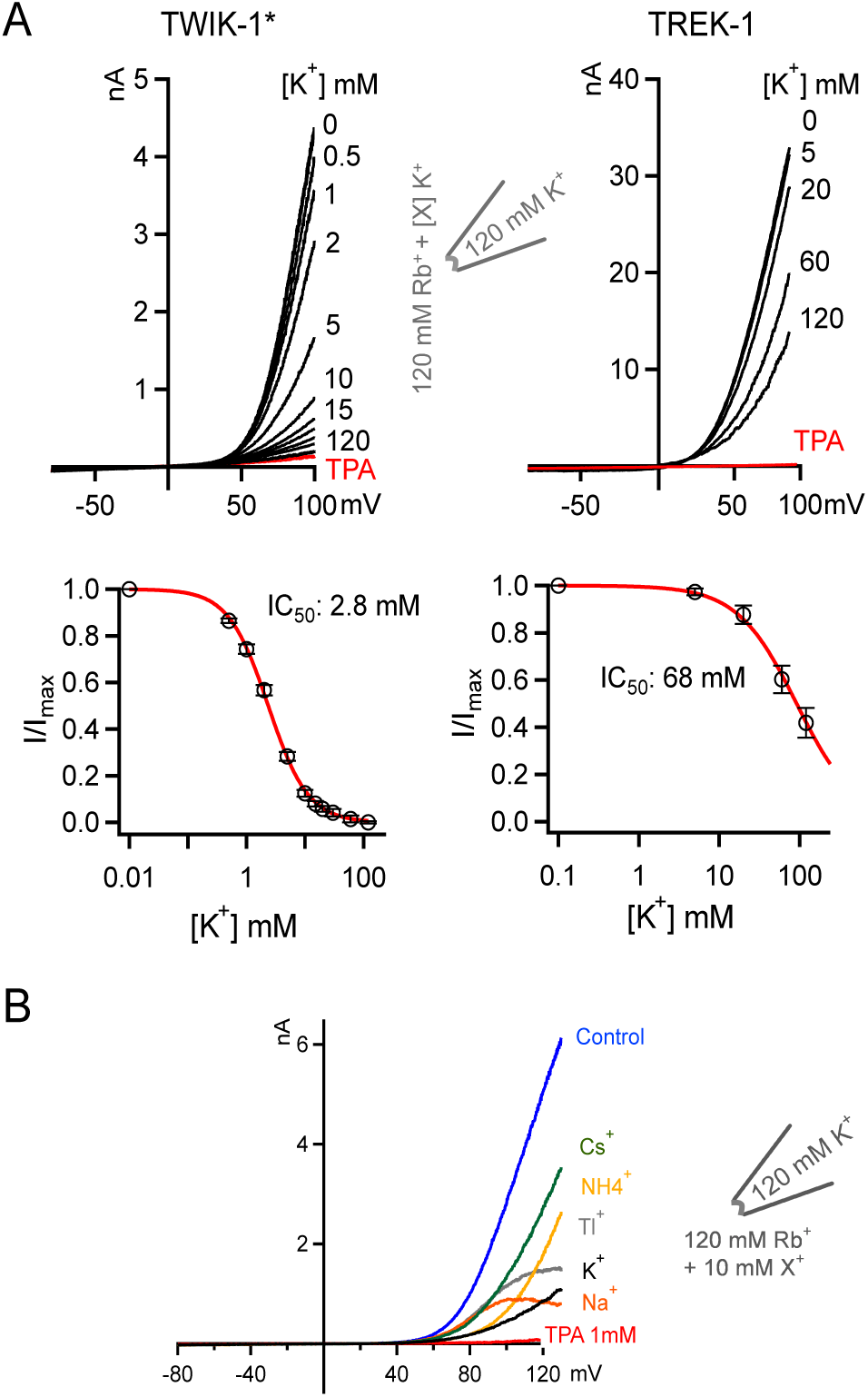
High-affinity inhibition of TWIK-1* Rb^+^ currents by intracellular K^+^. **A**) Comparison of K^+^ inhibition of Rb^+^ currents in TWIK-1* and TREK-1. Upper panel; in both cases the intracellular concentration of Rb^+^ is kept constant at 120 mM and increasing concentrations of K^+^ are added. Currents were recorded in response to a voltage ramp from -80 to +100 mV. In all cases these currents can be blocked by intracellular application of 1mM TPA (example shown in red). Lower panel; TWIK-1* currents from the experiments in the upper panel are plotted against intracellular K^+^ concentrations. **B**) Comparison of the inhibitory effect of different monovalent cations on TWIK-1* Rb^+^ currents. Currents were elicited in response to voltage ramps from -80 to + 120 mV. K^+^ is the most potent inhibitory ion within the physiological voltage range. Note that Cs^+^ which usually acts as a permeant blocker of most K^+^ channels has the least inhibitory effect.

### Contribution of the selectivity filter gate

We next set out to determine the mechanisms underlying this K^+^-dependent block of Rb^+^ permeation. In several previous studies we have shown that most forms of gating in K2P channels occur primarily within the SF (21,23-25,27). Interestingly, the SF of TWIK-1 exhibits several differences to other K2P channels within the canonical filter sequence as well as a dynamic K^+^>Na^+^ selectivity that can be modulated by mutations close to the filter (15). In that study it was reported that the T118I mutation in the first P-loop (T<u>T</u>GYG) ‘stabilized’ the channel in its K^+^-selective conformation, whilst the L108F/F109Y mutation in the preceding pore-helix stabilized its Na^+^-selective conformation.

We therefore examined the effect of these two mutations on the K^+^-dependent inhibition of WT TWIK-1* Rb^+^ currents. To perform these experiments, we expressed TWIK-1* in mammalian HEK293 cells and measured inward currents in the whole-cell configuration. Because Rb^+^ also activates the SF gate from the extracellular side (**Fig. 2C**) we measured the inhibitory effect of extracellular K^+^ on these inwardly directed Rb^+^ currents. **Fig. 5A** shows that in this recording configuration, WT TWIK-1* also still exhibited clear K^+^-dependent inhibition of these inward Rb^+^ currents, albeit with a reduced affinity (IC_50_ ∼ 22 mM).

**Figure 5.**
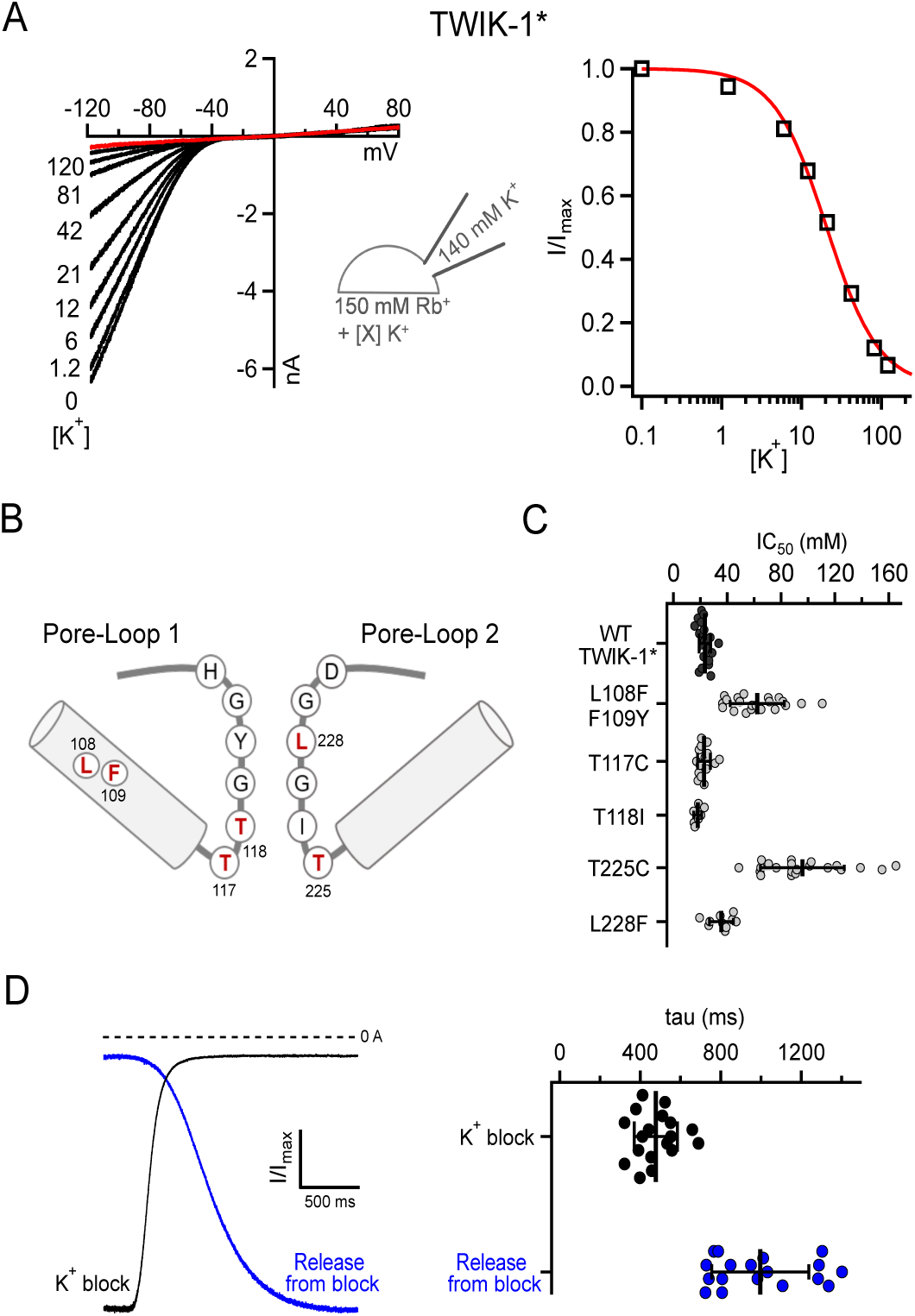
Inhibitory effect of extracellular K^+^ on whole-cell Rb^+^ currents. **A)** Inhibition of WT TWIK-1* by increasing concentrations of extracellular K^+^. Channels are expressed in HEK293 cells and the extracellular Rb^+^ concentration was kept constant (150 mM) with increasing concentrations of K^+^ as indicated. The right hand panel shows an example dose-response curve determined from the adjacent traces. **B)** A structural cartoon depicting the two pore-loops of TWIK-1 and the relative position of the mutations with the SF. **C)** Scatter plots of individual IC_50_ values determined from independent experiments similar to those shown in panel A. Data is shown for WT TWIK-1* and for the T118I, L108F/F109Y and L228F mutants which stabilize the dynamic K^+^/Na^+^ selectivity of TWIK-1 (15). Similar data is also shown for mutation of the two conserved threonines in the canonical K^+^-selective filter sequences of the first and second P-loops. The errors bars shown represent the mean and standard deviation. **D**) The difference in the time course of K^+^ block and release from K^+^ block is consistent with tight binding of K^+^ within the pore. Representative traces shows the rate of K^+^ block (black) and release from block (blue) as K^+^ or Rb^+^ solutions are applied, respectively. The right hand panel shows the values recorded from multiple individual experiments (see also Supplemental Fig. S3).

To examine the role of the SF, we studied the effect of K^+^ block on several mutants within the SF of TWIK-1* (**Fig. 5B**). The T118I mutation in TWIK-1* has previously been shown to stabilize the filter in an apparently K^+^-selective conformation (**Fig. 5B**). Consistent with this, we found that this mutant exhibited the same level of K^+^-dependent block as WT TWIK-1*, as did the L228F mutation which restores the consensus sequence of the second P-loop (**Fig. 5C**) and which also does not affect TWIK-1* K^+^-selectivity (15). However, the L108F/F109Y mutation within Pore-Helix 1 (**Fig. 5B**), which has been reported to reduce K^+^ selectivity (15), markedly reduced this inhibitory effect (IC_50_ ∼ 62 mM) (**Fig. 5C**).

### Functional asymmetry within the filter

Overall, these results strongly suggest a direct interaction of K^+^ with the SF of TWIK-1 that induces an unstable and therefore poorly conductive conformation of the SF gate. Interestingly, a recent study suggested that K^+^ exhibits a relatively low affinity for TWIK-1 (28). However, TWIK-1 activity was not measured directly and ion-exchange-induced difference IR spectroscopy of purified TWIK-1 channels in the absence of any transmembrane electrochemical potential was used to determine this interaction. By contrast, our approach provides a more direct measurement of how different ions permeate through the SF. Nevertheless, to further support our observations we examined the role of the filter gate in greater detail.

In a previous study of K2P channel gating, we have shown that, like KcsA, mutations within the conserved threonines of the consensus TxGYG sequence can have a profound effect on the SF gating mechanism (23). These conserved threonines are located at the lower entrance to the filter with their hydroxyl groups partially substituting for the waters of hydration of the permeant ion (usually K^+^) immediately prior to the ion entering the SF. We therefore mutated these two threonine residues and examined how they influence the high-affinity K^+^ block of Rb^+^ currents through TWIK-1* channels.

We found that mutation of the first conserved threonine (Thr^117^) had no effect (**Fig. 5C**), whereas mutation of Thr^225^ in the second P-loop markedly reduced this K^+^-dependent inhibition. This result not only supports the role of the SF in this interaction of K^+^ with TWIK-1, but also highlights the functionally asymmetric contribution of these two pore-forming loops to the SF gate.

To further support this inhibitory effect of K^+^, we also examined the kinetics of inhibition on WT TWIK-1* channels. Although the absolute values determined depend upon the limitations of the perfusion system, we found that the relative rates of block and unblock were different. Consistent with a tight binding of K^+^ within the pore, the rate of K^+^ inhibition was faster (τ ∼450 ms) than recovery of these currents upon washout (τ ∼ 1 s) (**Fig 5D** and **supplemental Fig S3**).

### Differential occupancy of K^+^ and Rb^+^ in the SF

To gain further insight into the mechanism by which K^+^ and Rb^+^ may interact differently within the filter of TWIK-1 to influence the SF gate, we employed atomistic double-bilayer molecular dynamics (MD) simulations of permeation similar to those we have recently used to examine permeation in other types of K^+^ channel, including K2P channels (23,24,29). Simulations with either K^+^ or Rb^+^ as the permeant ion were conducted (**Fig. 6**). This shows that over a total 6 µs simulation of outward permeation, the four main ion binding sites (S1-S4) within the filter of TWIK-1 display a markedly different occupancy by K^+^ compared to Rb^+^. In particular, whereas K^+^ exhibits preferential occupancy of the S2 and S4 sites, and relatively little occupancy at S1 and S3, Rb^+^ occupies all four sites, including S1 and S3. These results support previous studies which show K^+^ and Rb^+^ interact differently with the SF in other K^+^ channels (23,30) and suggest that a differential interaction of permeant ions at the S1 and S3 sites within TWIK-1 may contribute to its unusual functional properties.

**Figure 6.**
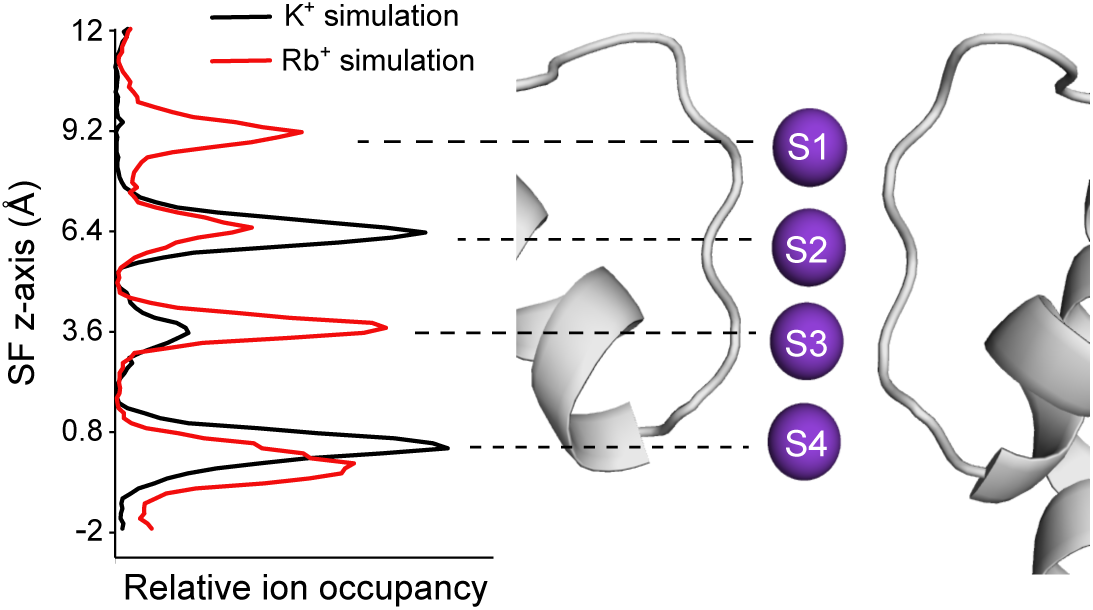
MD simulation reveals differential occupancy of SF by K^+^ vs Rb^+^. Double-bilayer simulations of TWIK-1 with either K^+^ or Rb^+^ reveals a different occupancy of the S1-S4 ion binding sites. The relative ion occupancy for outward currents are shown plotted against the position of the SF on the Z-axis (K^+^ in black; Rb^+^ in red). The data shown are collected from multiple repeats that each represent a total of 6 µs simulation during which 7 outward permeation events were observed for K^+^ and 13 event for Rb^+^. The peaks shown on the left correspond to the S1-S4 ion binding sites. These are structurally aligned with a depiction of the selectivity filter and pore-helices of TWIK-1.

## Discussion

Overall our results suggest that when present in the plasma membrane under normal physiological conditions and voltages, K^+^ stabilizes the TWIK-1 SF gate in an unstable, poorly conductive conformation. This is also supported by most functional studies that have attempted to record the functional activity of TWIK-1 using K^+^ as the permeant ion. Nevertheless, several previous studies have shown that the levels of whole-cell TWIK-1 channel K^+^ current can be enhanced by removal of an endocytic retrieval motif within the C-terminus (i.e. the TWIK-1* mutant used in this study) and our results are consistent with this observation, though these effects appear to be via changes in cell-surface expression rather than altering the gating properties of the channel *per se*. Other studies have also shown modest increases in whole-cell K^+^ currents of both WT TWIK-1 and TWIK-1* when Lys^274^ within the C-helix is mutated. Again, our results are also consistent with this observation, although the underlying mechanism still remains controversial. Some studies have proposed that post-translational sumoylation of Lys^274^ somehow silences channel activity, while other studies suggest that sumoylation is not responsible. Our results show that mutation of other charged residues within the C-helix can also produce similar increases in TWIK-1* whole-cell current, thus suggesting the effect of mutating Lys^274^ may not be as unique as previously thought. Either way, the increases in current seen with these C-helix mutations are relatively small in comparison to mutations within the ‘hydrophobic cuff’ of the inner cavity (e.g. L146N). However, even the effect of the L146N mutation is small in comparison to the large and robust currents that can be obtained when Rb^+^ is used as the permeant ion. This suggests that the unusual properties of the SF gate dominate the low intrinsic open probability of TWIK-1 when the channel is present in the plasma membrane.

In support of this we show that, like other K2Ps, TWIK-1 does indeed possess a flux-gating mechanism, but that it handles K^+^ and Rb^+^ differently and exhibits significantly altered properties to most other K2P channels. In particular, it displays a different ion activation profile, and with K^+^ this voltage-dependence is shifted into the supra-physiological range. Another major difference is that the SF gate can be activated by extracellular Rb^+^ and exhibits a highly unusual K^+^-dependent block. Our results demonstrate these unusual properties are conferred by the SF and that the contribution of the two pore-forming loops is unequal. This may be the result of an asymmetric contribution of the two P-loops to K2P channel gating and/or the unusual sequence of the consensus K^+^-filter motifs found in TWIK-1.

The physiological significance of these biophysical differences is unclear. One possibility may be related to the dynamic nature of TWIK-1 ion selectivity and that significant levels of TWIK-1 channels have been reported within endosomes where the unusual ionic environment and pH may modulate its functional properties (15,18,31). Also, the K^+^ concentrations within intracellular endosomes have been reported to be highly dynamic and proposed to play a role in the control of viral trafficking (32); in addition K2P channels are thought to play a key role in this process, though a specific role for TWIK-1 remains to be determined (33). Nevertheless, the unique biophysical properties we have identified in this study will clearly aid future functional and cellular studies of this enigmatic ion channel.

## Experimental Procedures

Electrophysiological studies of TWIK-1 and TREK-1 were performed as previously described (12,15). Briefly, human TWIK-1 was subcloned into the pFAW for expression oocytes and HEK-293 cells. Two-electrode voltage-clamp recordings of whole cell currents in *Xenopus* oocytes were recorded in ND96 solution at pH 7.4 (in mM, 96 NaCl, 2 KCl, 2 MgCl_2_, 1.8 CaCl_2_, 5 HEPES). For excised patch recordings, the intracellular solution (K^+^_int_) had the following composition in mM: 120 KCl, 10 HEPES, 2 EGTA, adjusted to the appropriate pH with HCl/KOH. The extracellular solution (K^+^_ex_, pipette solution) comprised, in mM; 120 KCl, 10 HEPES and 3.6 CaCl_2_ adjusted to pH 7.4 by HCl/KOH. In relevant recordings, the extra- and intracellular K^+^ was replaced by Cs^+^, Na^+^, NH_4_^+^, Rb^+^ or Tl^+^ and pH was adjusted by hydroxide of the relevant ion species and HCl. For recordings in HEK293 cells solutions had the following composition in mM; 150 KCl, 10 HEPES for extracellular solutions and pH was adjusted to 7.4 by HCl/KOH. Rb^+^ replaced K^+^ in other solutions and pH was adjusted by RbOH/HCl. Intracellular solutions were composed of 140 KCl, 10 HEPES and 2 EGTA. pH was adjusted to 8.0. All solution exchanges were done using a gravity driven multi-barrel micropipette.

For expression in oocytes, 50 nl of mRNA (500 ng/µl) was manually injected into Dumont stage VI oocytes and incubated at 17°C for 1-3 days prior to use. HEK293 cells were cultivated in DME-F12 Medium (Dulbecco’s Modified Eagle’s Medium and Ham’s F-12 Nutrient Mixture) and seeded in 5% gelatin treated Petri dishes 24 hours prior to the transfection. The transfection was done using FuGENE® HD Transfection Reagent according to the manufacturer’s protocol. Recordings were made 24-48 hours after transfection.

For excised patches from oocytes, pipettes were made using thick-walled borosilicate glass with a resistance of 0.3-0.9 MΩ and filled with the extracellular solutions. HEK293 cell experiments were done in whole cell configuration using extra thin glass pipettes with a resistance of 1-1.9 MΩ, filled with intracellular solutions. All pipettes were purchased from Science Products GmbH (Germany). Currents were recorded with an EPC10 amplifier running under Patchmaster software (HEKA electronics, Lamprecht, Germany) for excised patches, or Axopatch™ 200B running under Clampex (Molecular Devices, USA) and sampled at 10 KHz and 250, kHz respectively with analog filter set to 3 kHz and 10 KHz, respectively, and stored for later analysis. IC_50_ values were calculated by fitting data with the Hill-Equation: Base + (max – base)/[1 + (xhalf/x)h] where Base = zero current, max = maximum current, x = ligand concentration, xhalf = concentration for half-maximal occupation of the ligand binding site (inhibition by TPA or H^+^), rate = Hill coefficient. The voltage activated traces were fitted by the exponential function; y_0_ + A exp[-(t-t_0_)/*τ*], where y_0_ is the current where the fitting starts, A, the fit constant, t_0_ time where the fitting starts and *τ* time constant for the current increase. The time constant for block or release form the block are calculated by fitting the traces with the sigmoidal function; F(t)=A_base_+{(A_max_-A_base_)/[1+(t_1/2_/t)^n^]}, where A_base_ is the base current, A_max_ is the maximum current, t is the time and t_1/2_ is the time where the current reaches half the maximum current (A_max_) and n is the rise rate. Examples of the fits used are shown in the supplemental information. Unless otherwise described, values result from at least three independent experiments given as average ± standard error of the mean (SEM).

MD simulations were performed as described before (23,24,29,34). Briefly; we used a modified version of GROMACS 4.6, the AMBER99sb force field and the SPC/E water model for the equilibrium and production simulations. The simulations were performed at 300 K with a velocity rescaling thermostat and the pressure kept at 1 bar using a semi-isotropic Berendsen barostat. All bonds were constrained with the LINCS algorithm and using virtual sites for hydrogen atoms allowed simulations to be performed with a 4 fs integration time step. The crystallographic structure of TWIK-1 (PDB ID code: 3ukm (9) was adopted as the initial structure in the MD simulations and was embedded into an aqueous POPC lipid bilayer with 0.6 M KCl. The system was equilibrated for 20 ns with position restraints on all heavy atoms using a force constant of 1000 kJ mol^-1^ nm^-2^ to the reference structure, followed by an additional 10 ns simulation without position restraints. Production runs were then performed on the equilibrated TWIK-1 structure using a computational electrophysiology protocol (34). For K^+^ and Rb^+^, 20 replicas of 300 ns were carried out for each ion type, respectively. Potential gradients were generated by introducing a charge difference of 2 K^+^ or 2 Rb^+^ ions between the two compartments separated by the two lipid bilayers. During the MD simulations, the number of the ions was kept constant by an additional algorithm (34). The potential difference between two compartments corresponds to 360 ± 70 mV and 390 ± 70 mV for K^+^ and Rb^+^ simulations, respectively.

## Acknowledgements

This work was supported by grants from the Biotechnology and Biological Sciences Research Council (BBSRC), the Deutsche Forschungsgemeinschaft (DFG, Research Group FOR 2518 DynIon) and the Centre National de la Recherche Scientifique (CNRS). F.A-W was supported by a King’s Scholarship from the Malaysian Government.

## Supporting Information

**Supplemental Fig. S1.**
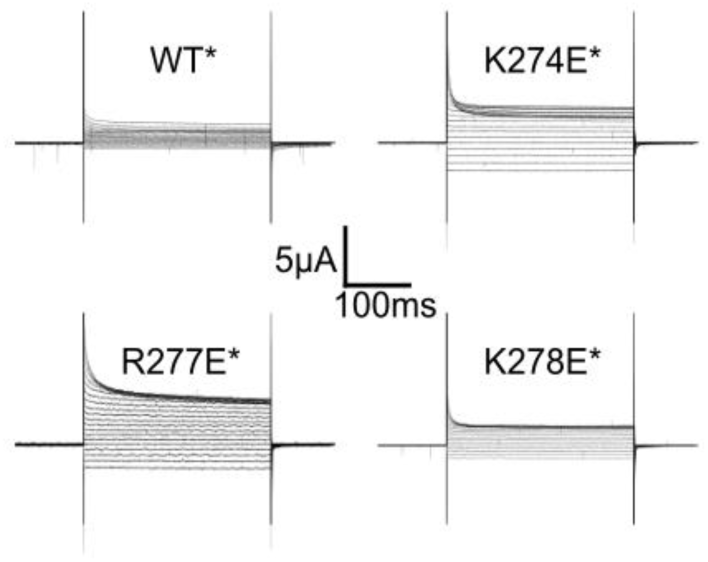
Example currents for WT TWIK-1* and the three key mutations shown in Figure 1B. These mutants all produce increases in whole-cell current that are equivalent to the effect of the previously reported K274E mutation (8). Currents were recorded by TEVC in response to voltage steps from -120 to +60 mV from a holding potential of -80 mV. Reported values were measured at 0 mV.

**Supplemental Fig. S2.**
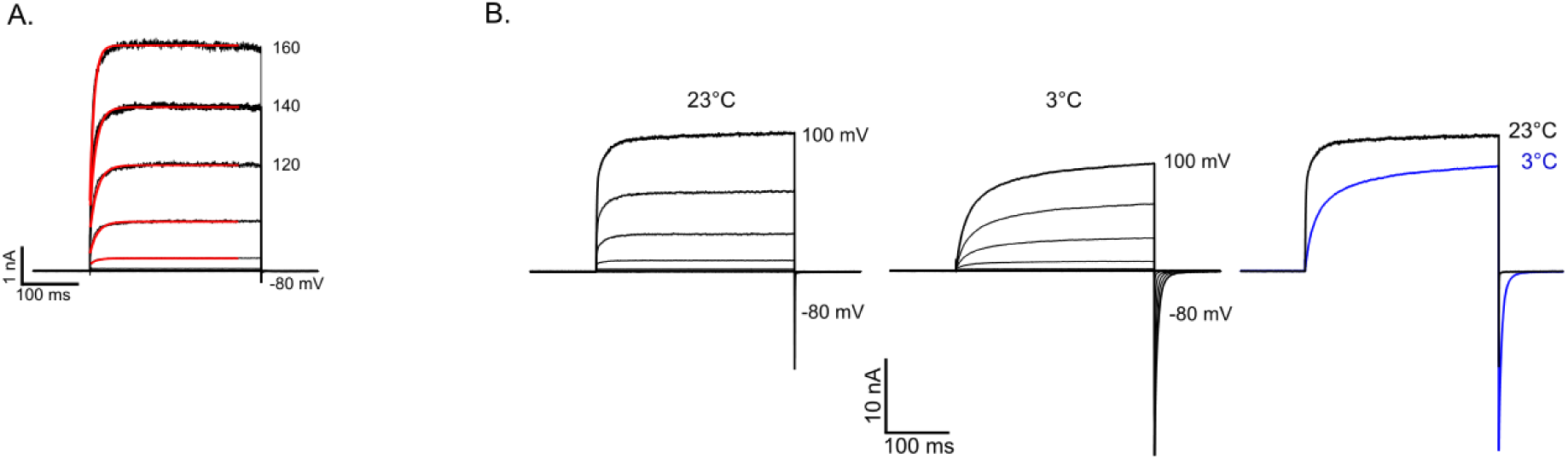
**A)** Example of the fits used (red) on Rb^+^ activated TWIK-1 currents. These fits were used to obtain the rate constants shown in Fig.3C which demonstrate that activation is not voltage-dependent (see methods for details). **B)** Currents recorded in excised patches from a mutant version of TREK-2 (Gly^67^-Glu^340^) (26) at both 23° and 3° under the same conditions shown in Fig. 3D. At 23° no tail currents are visible, but upon cooling to 3° the rate of activation slows, as well as the tail currents so they become more visible. An overlay of these currents at +100 mV is shown in the right hand panel.

**Supplemental Fig. S3.**
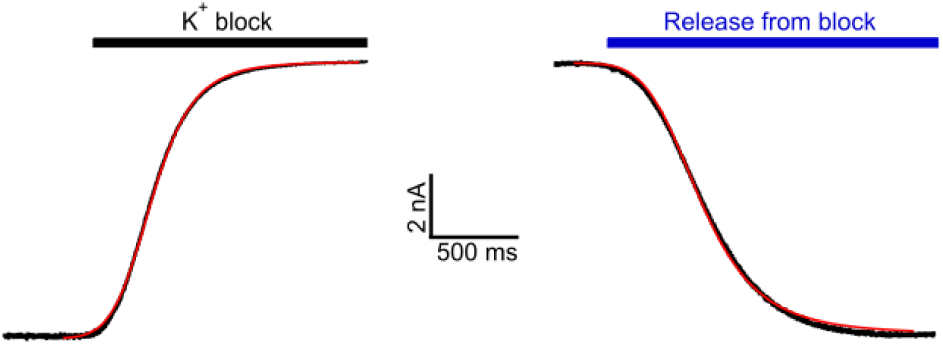
Example of the fits used (in red) to obtain the rate constants for K^+^ block and unblock that are shown in Fig. 5D. See experimental procedures for details.

